# Dentate gyrus and CA3 activity mediates light-tone second-order conditioning expression in mice

**DOI:** 10.1101/2025.08.29.673009

**Authors:** Marc Canela, Jordi Bach Adell, David Roura Coll, Jose Antonio González-Parra, Julia S. Pinho, Arnau Busquets-García

## Abstract

Second-order conditioning (SOC) enables animals to form complex predictions about their environment, even in the absence of direct experience. While the neural mechanisms underlying first-order conditioning (FOC) are well characterized, the circuits supporting SOC expression remain poorly understood. To address this gap, we investigated the brain regions and cell types involved in SOC recall in mice and tackled the technical challenges of quantifying brain-wide neural activity. We employed a light–tone SOC paradigm in TRAP2:Ai14 mice, which allowed us to tag neurons active during SOC recall via tdTomato expression. Applying generalized linear models, we identified that the activity in the dentate gyrus (DG) and CA3 regions of the dorsal hippocampus significantly associated with SOC-related behavioral responses. To test their functional relevance, we used chemogenetic inhibition of CaMKII^+^ neurons in these regions, which confirmed a causal role for DG/CA3 circuits in SOC recall. Together, our results highlight the dorsal hippocampus as a critical substrate for retrieving indirectly learned associations.

## INTRODUCTION

The brain’s ability to encode, store, and recall memories^1^ remains one of the least understood parts of human biology. Indeed, the brain works differently depending on the type of memory, such as episodic, procedural, semantic, and others^2,3^. Among these, a large portion is unconscious, mainly operating through associative mechanisms^4,5^. In such cases, a specific cue, like a smell, sound, or place, can bring up the memory of another stimulus or trigger a behavioral response^6^.

These associative processes include various subtypes corresponding to different conditioning paradigms^4^. In first-order conditioning (FOC), an animal learns to associate a neutral stimulus, such as a light or tone, with a biologically significant unconditioned stimulus (US), like food or an electric footshock. After repeated pairings, the neutral cue becomes a conditioned stimulus (CS) and is enough to trigger alone a conditioned response similar to the US^7,8^.

In contrast, higher-order conditioning forms an indirect link between a specific CS and the US, rather than a direct association. This type of learning includes two primary forms: sensory preconditioning (SPC) and second-order conditioning (SOC). In SPC, the process begins by pairing two neutral stimuli, such as a light and a tone. After the animal establishes an association between them, one of those stimuli is then associated with the US. As a result, the cue directly paired with the US becomes the first-order conditioned stimulus (CS_1_), while the other becomes the second-order conditioned stimulus (CS_2_). In SOC, the sequence reverses: a neutral stimulus first pairs with the US, then this CS_1_ links to another neutral stimulus. Remarkably, in both SPC and SOC, CS_2_ can trigger conditioned responses, even though CS_2_ lacks a direct association with the US^5,9,10^. Therefore, this ability to increase the set of stimuli an animal can respond to makes higher-order learning crucial for survival^10,11^. However, when these processes become disrupted, they may contribute to disorders where individuals maintain persistent fear responses through mechanisms like SOC^12,13^. Thus, understanding how the brain encodes and retrieves such complex associations is vital for explaining both adaptive behavior and the neural basis of cognitive disorders.

Although researchers have well characterized the neural basis of FOC^7^, the brain mechanisms supporting SPC and SOC remain less understood. In fact, there is more research on SPC than SOC, mainly because it addresses how the brain links two initially neutral stimuli, an associative process that FOC and SOC fail to tackle. As a result, numerous investigations have mapped some circuits involved in SPC, highlighting important regions such as the retrosplenial cortex (RSC)^14^, the orbitofrontal cortex (OFC)^5^, the perirhinal cortex (PRh)^15^, the dorsal hippocampus^16^, the amygdala, especially the basolateral amygdala (BLA)^17,18^, as well as the nucleus accumbens (NAc) and ventral tegmental area (VTA)^19^. On the other hand, some research has begun to show causal involvement of specific brain regions involved in SOC, including the BLA^18,20^, the lateral hypothalamus (LH)^21^, and the parabrachial nucleus (PBN)^22^. However, few studies have investigated the brain substrates of SOC recall in mice and whether these circuits are shared with FOC expression.

To investigate the brain circuits underlying associative learning and other behaviors, researchers often analyze fluorescent activity markers in brain slices^23–26^. However, manual analysis of fluorescence images is slow, prone to error, and susceptible to observer bias and fatigue. These factors reduce reproducibility and make manual methods less reliable for large-scale studies. On the other hand, automated tools provide better accuracy, consistency, and scalability^27,28^. Yet, they also come with significant limitations: many require extensive customization to fit different experimental protocols and tissue types, demand high computational power, and rely heavily on labor-intensive image labeling for training. Additionally, pre-trained models often struggle to generalize across different datasets, which creates inefficiencies that slow down research progress.

In this study, we address the limitations of previous approaches by integrating advanced behavioral, imaging, and computational techniques to investigate the brain substrates underlying recall in SOC. Using TRAP2:Ai14 mice, a proxy of cell activity using the *Fos* promoter^29^, we genetically tagged neurons activated during the recall of CS_1_ and CS_2_ in a light-tone SOC paradigm previously validated in our laboratory^30^. We conducted a behavioral analysis using DeepLabCut^31^, Keypoint-MoSeq^32^, and DeepOF^33^, enabling the extraction of robust metrics for modeling brain activity. To overcome the constraints of manual histological analysis, we developed CellRake, an open-source Python package for automated and accurate quantification of fluorescent markers in brain slices. This tool facilitated the identification of glutamatergic neurons in the dentate gyrus (DG) and CA3 hippocampal subregions, which are activated during CS_2_ recall and are associated with fear-related behaviors. To assess the causal role of these excitatory hippocampal neurons, we employed chemogenetic inhibition during SOC recall. Together, this multimodal approach identified CaMKII^+^ neurons in the DG/CA3 as key contributors to CS_2_ recall in SOC expression.

## RESULTS

### Second-order conditioning expression in TRAP2:Ai14 mice

To investigate the neural substrates underlying the recall of first- and second-order conditioning, we used TRAP2:Ai14 mice, which enable the permanent genetic labeling of neurons activated during a specific time window, via CreER expression driven by the *Fos* promoter.^29^. This method enabled us to selectively tag neurons involved in recalling conditioned stimuli, specifically a tone (CS_2_) and a light (CS_1_). We employed a previously validated SOC protocol in our laboratory^30^, consisting of five stages on separate days: habituation, pairing a CS_1_ (e.g., a light) with a foot shock, pairing a CS_2_ (e.g., a tone) with CS_1_, and conducting two independent probe tests for both conditioned stimuli. To label neurons active during the CS_2_ recall, we administered 4-hydroxytamoxifen (4-OHT) immediately after this probe session, which triggered tdTomato expression in active neurons exposed to CS_2_. One week later, after enough time for tdTomato to express, we performed the CS_1_ recall test and collected the brains for cFos immunohistochemistry, allowing us to identify neurons active during the CS_1_ late recall. This experimental design enabled us to map and quantify brain-wide activity patterns associated with each stage of SOC and relate them to behavioral responses (Figure 1A).

**Figure 1.**
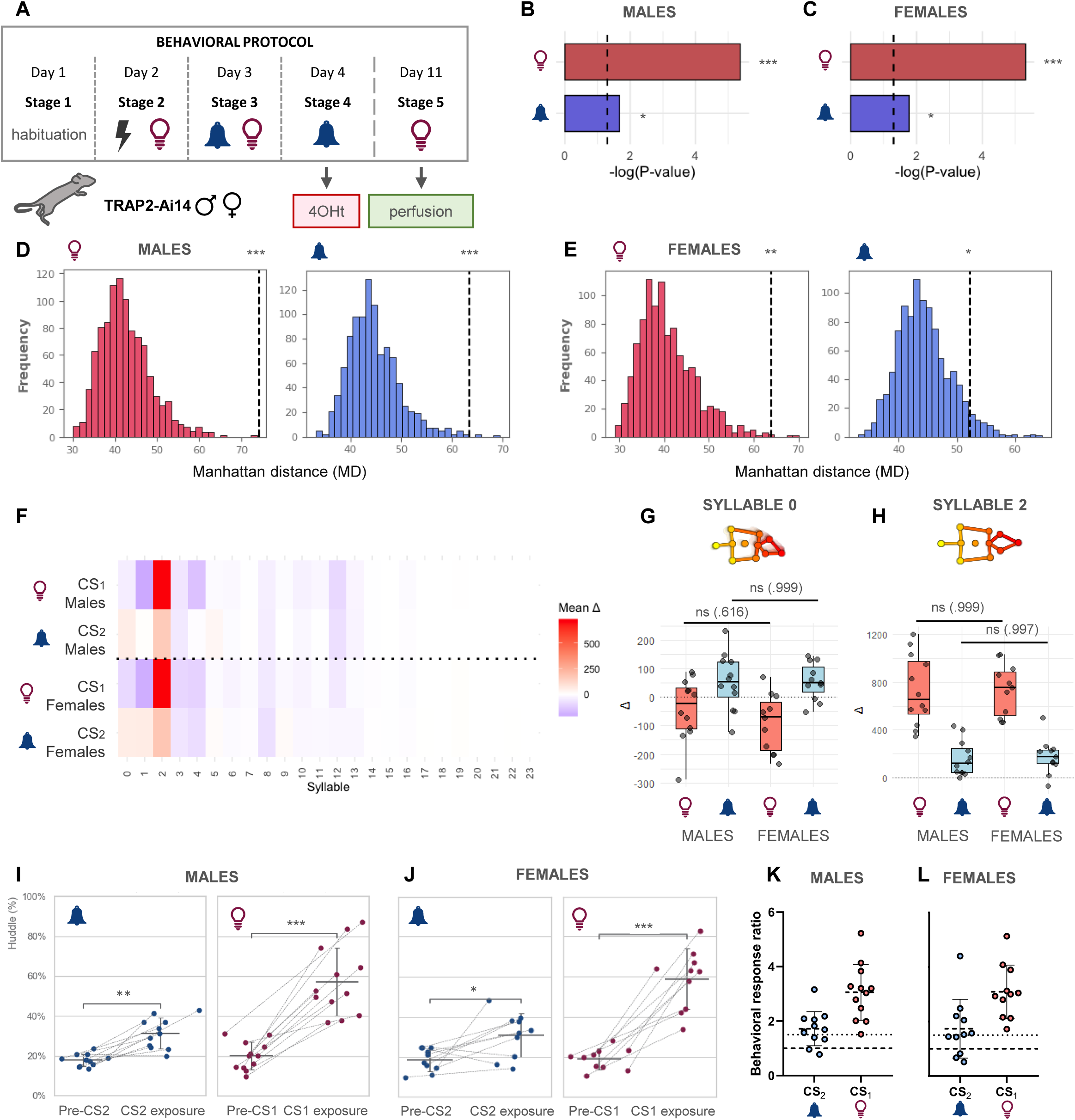
Second-order conditioning recall induces robust behavioral chang in TRAP2:Ai14 mice. (A) Schematic representation of the light-tone behavioral SOC protocol using TRAP2:Ai14 animals. Mice underwent habituation (Stage 1), first-order conditioning with light and footshock (Stage 2), second-order conditioning with tone paired with light (CS_1_) (Stage 3), and independent recall sessions for tone (CS_2_) (Stage 4) and light (CS_1_) (Stage 5). We tagged active neurons during CS_2_ recall using 4-hydroxytamoxifen (4-OHT), waited one week, and collected the brains after the CS_1_ recall. (B–C) Outcomes from Hotelling’s T^2^ analysis on the Δ metrics, shown as the negative logarithm of the p-value for each tested experimental condition: CS_1_ males (p *<*0.0001, n = 12), CS_2_ males (p = 0.0210, n = 12), CS_1_ females (p *<*0.0001, n = 11), and CS_2_ females (p = 0.0167, n = 11). (D–E) Transition dynamics measured through the Manhattan distance (MD) across conditions: CS_1_ males (p *<*0.0001, n = 12), CS_2_ males (p = 0.000114, n = 12), CS_1_ females (p = 0.000162, n = 11), and CS_2_ females (p = 0.0424, n = 11). (F) Heatmap of average syllable Δ vectors showing up-regulation (red) or down-regulation (blue) in syllable abundance. (G) Differences of average Δ values in syllable 0 between sexes: CS_1_ (p = 0.616, males: n = 12, females: n = 11), CS_2_ (p = 0.999, males: n = 12, females: n = 11). (H) Differences of average Δ values in syllable 2 between sexes: CS_1_ (p = 0.999, males: n = 12, females: n = 11), CS_2_ (p = 0.997, males: n = 12, females: n = 11). (I–J) Huddle percentage computed through DeepOF across conditions: CS_2_ males (p = 0.00195, n = 12), CS_1_ males (p = 0.0005, n = 12), CS_2_ females (p = 0.0167, n = 11), and CS_1_ females (p *<*0.0001, n = 11). (K–L) Behavioral response ratios (post/pre-immobility). The darker line indicates a ratio of 1, corresponding to the brink between increase and decrease. We established a threshold of 1.5 (thinner line) to classify animals as high- or low-performers for subsequent neural analyses. P-values are denoted as p *<*0.05 (*), p *<*0.01 (**), and p *<*0.001 (***). The data is presented as mean ± SD.

For this experiment, we used adult male and female mice to replicate the behavioral findings reported in our recent work^30^. We estimated animal pose using DeepLabCut^31^, and analyzed the resulting trajectories with the unsupervised Keypoint-MoSeq algorithm^32^, which decomposes behavior into stereotyped motifs known as behavioral syllables.

We first measured the amount of time each animal spent producing each syllable and calculated the change in time between the post-stimulus and pre-stimulus periods. These differences, called Δ scores, show whether a syllable’s usage increased (Δ *>*0) or decreased (Δ *<*0) after stimulus presentation. To determine if these changes indicated a real behavioral modulation, we used a multivariate Hotelling’s T^2^ test to see if the set of Δ values differed significantly from zero. In males (Figure 1B), the Δ values were markedly different from zero in response to CS_1_ (p *<*0.0001) and CS_2_ (p = 0.0210). Similarly, in females (Figure 1C), the Δ values also significantly differed from zero in response to CS_1_ (p *<*0.0001) and CS_2_ (p = 0.0167), showing that both conditioned cues caused consistent behavioral changes in both sexes.

As previously performed^30^, we then calculated transition matrices between syllables to examine the dynamics of behavior. We assessed the change by measuring the Manhattan Distance (MD) between different time points and derived an empirical p-value through bootstrapping. This dynamic analysis confirmed the findings from Hotelling’s T^2^ test. In males (Figure 1D), both CS_1_ (MD = 75, p *<*0.0001) and CS_2_ (MD = 65, p = 0.000114) showed significant shifts in their transitions. Similarly, females (Figure 1E) also exhibited a significant shift to CS_1_ (MD = 65, p = 0.000162) and CS_2_ (MD = 53, p = 0.0424).

We next aimed to identify a behavioral metric that we could correlate with neural activity. To this end, we analyzed the Δ values across syllables (Figure 1F). Our results replicated previous findings involving syllables 0 and 2^30^. Specifically, for syllable 0 (Figure 1G), which corresponds to walking, we observed a downregulation in response to CS_1_, with a mean Δ value of approximately –50 and no significant sex differences (p = 0.616). In contrast, the response to CS_2_ showed an upregulation, with a mean Δ value around +50, again without significant differences between sexes (p = 0.999). For syllable 2, representing immobility, both CS_1_ and CS_2_ evoked strong upregulation, with mean Δ values exceeding +700 for CS_1_ and +150 for CS_2_ (Figure 1H). These effects were consistent across sexes, with no significant differences for either CS_1_ (p = 0.999) or CS_2_ (p = 0.997).

While acknowledging the complexity of behavior and the detailed information obtained by each syllable, it is essential to select a metric that effectively summarizes behavioral change. Considering that the highest Δ scores for both CS_1_ and CS_2_ belong to syllable 2, representing immobility-like behaviors, we chose this type of response for downstream modeling with brain activity. Specifically, we focused on an immobility-related measure, huddling, which we can automatically quantify using DeepOF^33^, and we previously confirmed that it strongly correlates with freezing measurements^30^. The analysis of huddling displays significant changes in response to CS_1_ and CS_2_ in both males (Figure 1I) and females (Figure 1J).

Furthermore, a closer examination of individual values for our immobility measure revealed variability in performance. By calculating the behavioral response ratio between post-stimulus and pre-stimulus periods, we found that the lowest response ratio to CS_1_ was 1.5, indicating at least a 50% increase in immobility after the first-order stimulus. Based on this, we set a threshold of 1.5 for both CS_1_ and CS_2_ in males (Figure 1K) and females (Figure 1L) to classify animals as high- or low-performers in our second-order conditioning paradigm.

Overall, using TRAP2:Ai14 mice, we confirmed the effectiveness of our SOC protocol. Behavioral analysis with DeepLabCut and Keypoint-MoSeq showed significant and consistent changes in syllable expression after CS_1_ and CS_2_ presentations in both sexes. Notably, immobility-related behavior (syllable 2) exhibited strong upregulation, leading us to select immobility as a reliable behavioral measure for further analysis alongside neural activation.

### CellRake accurately quantifies fluorescent markers in histological brain sections

After establishing a behavioral metric to correlate behavioral responses and brain activity, we focused on quantifying this activity by counting tdTomato- and cFos-positive cells across various brain regions. Due to the large number of animals from both sexes and the involvement of multiple brain areas, manual counting would be tedious and prone to bias. To address this, we developed CellRake, a Python package that leverages several basic Python libraries, including NumPy^34^ for numerical operations, pandas^35^ for data manipulation, scikit-image^36^ for advanced image analysis techniques, scikit-learn^37^ for predictive data analysis, and Matplotlib^38^ for plotting. Only relying on these basic, well-known packages facilitates a robust environment that enhances CellRake’s usability, reliability, and maintainability.

CellRake uses a two-step workflow (segmentation followed by classification) to detect cells expressing positive fluorescent markers such as tdTomato and cFos (Figure 2A). To assess CellRake’s performance throughout this process, we used a benchmark dataset of 153 images of the basolateral amygdala (BLA) from 36 male mice, since this brain region is a key player in higher-order conditioning^17,18,39^ and its cellular architecture tends to be more uniform^40^. All images showed of cFos nuclear labeling and cytoplasmic tdTomato markers resulting from genetic recombination on TRAP2:Ai14.

**Figure 2.**
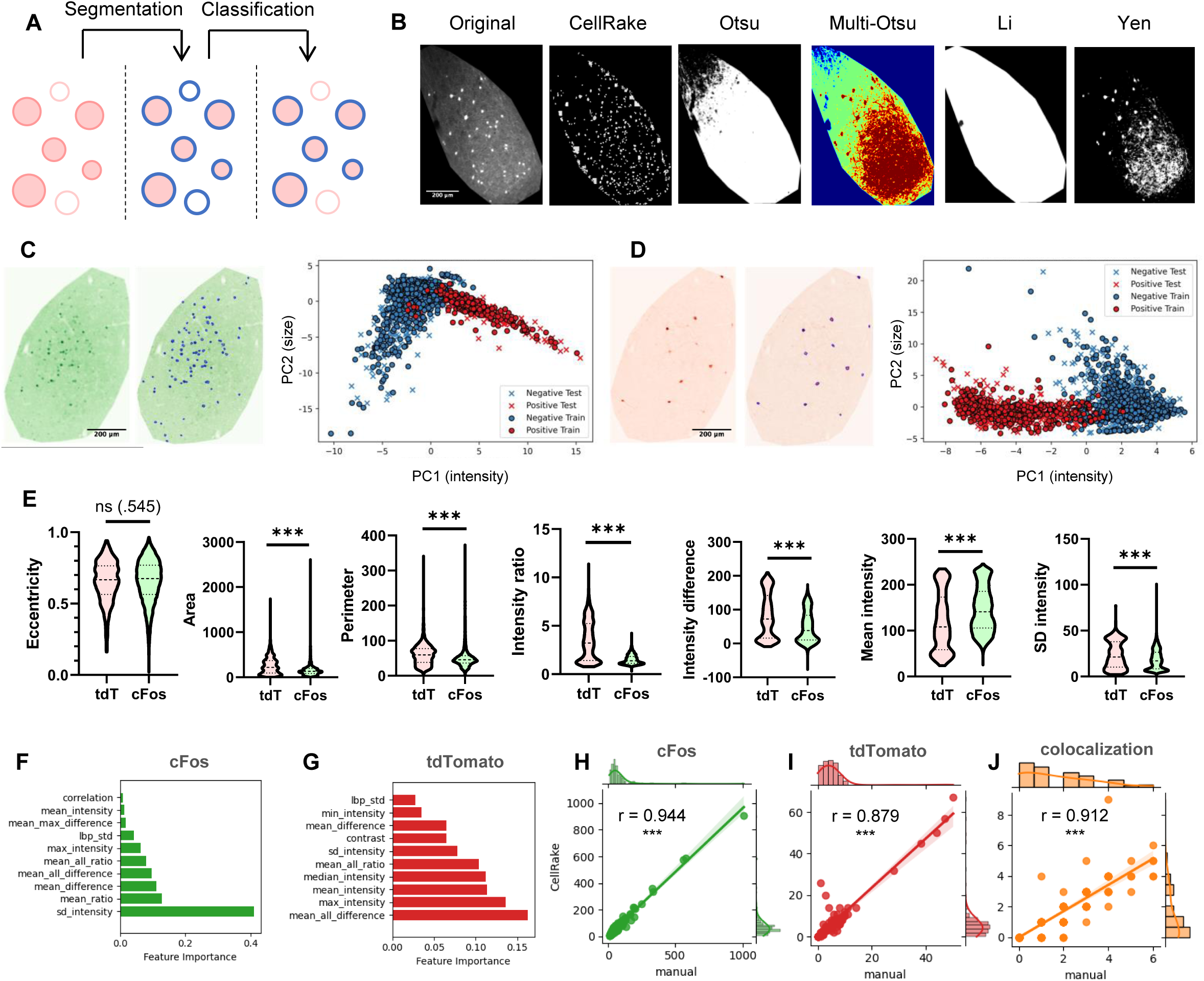
Overview and performance evaluation of CellRake’s cell detection pipeline. (A) Schematic representation of CellRake in two phases: segmentation of candidate blobs and machine learning–based classification to identify true positives. (B) Comparison of CellRake’s segmentation (second panel) against standard thresholding methods (Otsu, Multi-Otsu, Li, Yen), highlighting its ability to preserve candidate blobs across heterogeneous intensity backgrounds. (C-D) Classifier results for cFos- and tdTomato-labeled cells. Left: example of nuclear cFos and cytoplasmatic tdTomato staining, along with the identified true positive blobs. Right: PCA of 22 extracted features using two components: PC1 enriched in color intensity features and PC2 enriched in size features. The actual markers (in red) separate from the false-positive blobs (in blue). (E) Feature analysis comparing tdTomato^+^ (n = 990) and cFos^+^ (n = 12642) cells: eccentricity (p = 0.545), area (p *<*0.0001), perimeter (p *<*0.0001), intensity ratio (p *<*0.0001), intensity difference (p *<*0.0001), mean intensity (p *<*0.0001), SD intensity (p *<*0.0001). (F-G) Feature importance analysis in the cFos and tdTomato classifiers. CellRake mainly depends on intensity-based features, while size and shape play a minor role. (H–J) Correlation of CellRake’s automated counts with manual ground truth for cFos (r = 0.944, p *<*0.0001, n = 153), tdTomato (r = 0.879, p *<*0.0001, n = 153), and their colocalization (r = 0.912, p *<*0.0001, n = 153). P-values are denoted as p *<*0.05 (*), p *<*0.01 (**), and p *<*0.001 (***). The data is presented as mean ± SD.

In the initial segmentation phase, the algorithm processes single-layer images to extract structures resembling potential positive detections (i.e., blobs), favoring recall over precision. This strategy ensures that all possible positives (true positives and false positives) are included in the initial pool of candidates. Comparing CellRake’s segmentation method with other thresholding techniques, our approach preserves the integrity of detected regions by removing only the surrounding background of each blob, allowing for robust detection even in images with variable intensities or heterogeneous backgrounds. A visual inspection of images with non-uniform surroundings shows that CellRake consistently retains candidate blobs under challenging conditions. In contrast, traditional thresholding methods such as Otsu, Multi-Otsu, Li, and Yen often produce incomplete or fragmented segmentations (Figure 2B).

Once segmentation is complete, we independently trained two random forest classifiers, one for each marker (cFos and tdTomato). For each candidate blob, CellRake extracts 22 features, including metrics such as brightness, area, perimeter, and eccentricity. These features are then used as input to the machine learning classifier, which evaluates each candidate as either a true or false positive. Ultimately, CellRake retains only the blobs classified as true positives, achieving its goal of accurate and reliable cell detection.

For the cFos-labeled images, CellRake required 15 minutes to segment all the blobs. Then, on the training set, the classifier achieved a ROC AUC of 0.987, an average precision (AP) of 0.976, a precision of 0.971, a recall of 0.913, and an F1 score of 0.941. On the test set, the model exhibited strong generalization performance, with a ROC AUC of 0.993, AP of 0.979, precision of 0.963, recall of 0.883, and F1 score of 0.922 (Figure 2C).

In contrast, the segmentation of tdTomato-labeled images finished in just 7 minutes. The corresponding classifier achieved near-perfect metrics on the training set: a ROC AUC of 0.999, AP of 0.999, precision of 0.992, recall of 0.983, and an F1 score of 0.987. These high values remained on the test set, with a ROC AUC of 0.9998, AP of 0.9996, precision of 0.992, recall of 0.994, and F1 score of 0.993 (Figure 2D).

Feature analysis using CellRake confirmed that both markers share a similar elliptical shape (eccentricity = 0.65, p = 0.545), though tdTomato^+^ markers are 33.4% larger (p *<*0.0001) and have a 13% longer perimeter (p *<*0.0001). Regarding intensity, tdTomato markers display significantly higher contrast with a 34% greater mean intensity ratio (p *<*0.0001) and a 52% larger cell-to-background difference (p *<*0.0001). At the same time, cFos^+^ cells show 23% higher mean absolute intensity (p *<*0.0001) and reduced variability (p *<*0.0001) (Figure 2E).

Examining which features drive the classifier’s decision-making, we found that CellRake heavily prioritizes intensity-based features, regardless of the marker type. For cFos, it relies more on features such as the standard deviation of blob intensity and mean ratio, which better capture dynamic intensity patterns (Figure 2F). In contrast, for tdTomato, where the contrast with the background is high, features like the difference between blob intensity and background intensity, maximum intensity, and mean intensity dominate the decision-making process (Figure 2G). Interestingly, shape and size features, such as area, perimeter, and axis lengths, play a negligible role in both cases. This indicates that the classifier primarily focuses on intensity metrics to differentiate between true positives and background noise.

To evaluate the global performance of CellRake, we compared the classifier’s output counts for cFos and tdTomato with manual counts, as well as the colocalization output with manually assessed values. The results indicate that CellRake can detect and colocalize cFos^+^ and tdTomato^+^ cells in all three cases, with automatic counts closely matching ground truth values. For cFos, the mean classified positives (82.627) closely resemble the ground truth (79.725), with a strong and significant correlation (r = 0.944, p *<*0.0001) (Figure 2H). Similarly, for tdTomato, the average classified positives (6.471) closely align with manual counts (5.569), as evidenced by a strong and significant positive correlation (r = 0.879, p *<*0.0001) (Figure 2I). The colocalization analysis also achieves outstanding results, with a high and significant correlation (r = 0.912, p *<*0.0001) and the average number of colocalized cells identified by CellRake (1.431) closely approximating the manual value (1.699) (Figure 2J).

Overall, evaluating CellRake on a benchmark dataset of cFos- and tdTomato-labeled images, the package achieved high accuracy with near-perfect classification metrics, especially for tdTomato^+^ cells. We also observed that differences in marker properties influenced performance, with intensity features playing a key role in driving classification decisions. Importantly, CellRake’s automatic cell counts and colocalization results showed strong, significant correlations with manual counts, demonstrating its reliability and efficiency for large-scale marker quantification.

### DG and CA3 link behavioral performance to neural activation during secondorder memory recall

After benchmarking CellRake, we applied this software to measure the number of positive cells in different brain regions from the TRAP2:Ai14 mice shown in Figure 1. For each region, we analyzed between four and six brain slices per animal.

After obtaining cell counts for each slice, we then assessed the relationship between these neural activation profiles and behavioral performance using a linear model. Specifically, we fitted separate generalized linear mixed models (GLMMs) using immobility ratios (i.e., the ratio of huddling between pre- and post-presentation of CS_1_ or CS_2_). We used individual brain regions for each model, except for the DG and CA3, in which we combined them into one model with a fixed effect, given their close anatomical proximity and the fact that the DG encloses part of the CA3. Overall, these separated models enabled us to evaluate the contribution of each region independently, thereby preventing multicollinearity and enhancing model interpretability. We used the raw, unnormalized counts without averaging, but included the logarithm of the ROI area as an offset and the individual IDs as a random effect to account for repeated measures. We also included other fixed effects such as sex and the behavioral performance category (highvs. low-performers), this last only in the case of tdTomato count, since all animals were high-performers for cFos.

Initially, we fitted a GLMM assuming a Poisson distribution because it is suitable for discrete data, such as counts. However, the tdTomato and cFos data showed overdispersion, meaning the variance was more than twice the mean, which breaks the Poisson assumption that the mean equals the variance. To address this, we used a Negative Binomial distribution instead, as it handles overdispersion.

When fitting a GLMM for tdTomato labeling, which marks neurons activated during CS_2_ memory recall in TRAP2:Ai14 mice, we found evidence of zero inflation. In other words, the number of zero counts was higher than expected under standard count models, indicating an excess of brain slices with no tdTomato^+^ cells. To account for this, we incorporated different zero-inflation structures into the models to capture the likelihood of excess zeros. We then compared these models using the Akaike Information Criterion (AIC) to determine the best approach for handling zero inflation in our count data.

When comparing the effects of behavioral response ratios on tdTomato counts across brain regions, only the dorsal DG and CA3 showed a significant coefficient (p = 0.00389) (Figure 3A), with no significant differences across sex (p = 0.09). Between the two brain regions, DG exhibited significantly higher tdTomato^+^ cell counts than CA3, with more than twice the expected number of labeled cells (p *<*0.0001). Notably, the interaction between the immobility ratio and the behavioral performance category (high- and low-performers) was also significant (p = 0.0429), indicating that performance level differently influences the neural activation patterns associated with behavioral responses. For high performers, a one-unit increase in the response ratio yields a 35.5% increase in expected tdTomato^+^ cell counts, holding other variables constant. In contrast, for low performers, this effect not only disappeared but also slightly reversed, with a one-unit increase in the behavior ratio corresponding to a 28.2% decrease in expected cell counts, assuming all other variables unchanged (Figures 3B–C).

**Figure 3.**
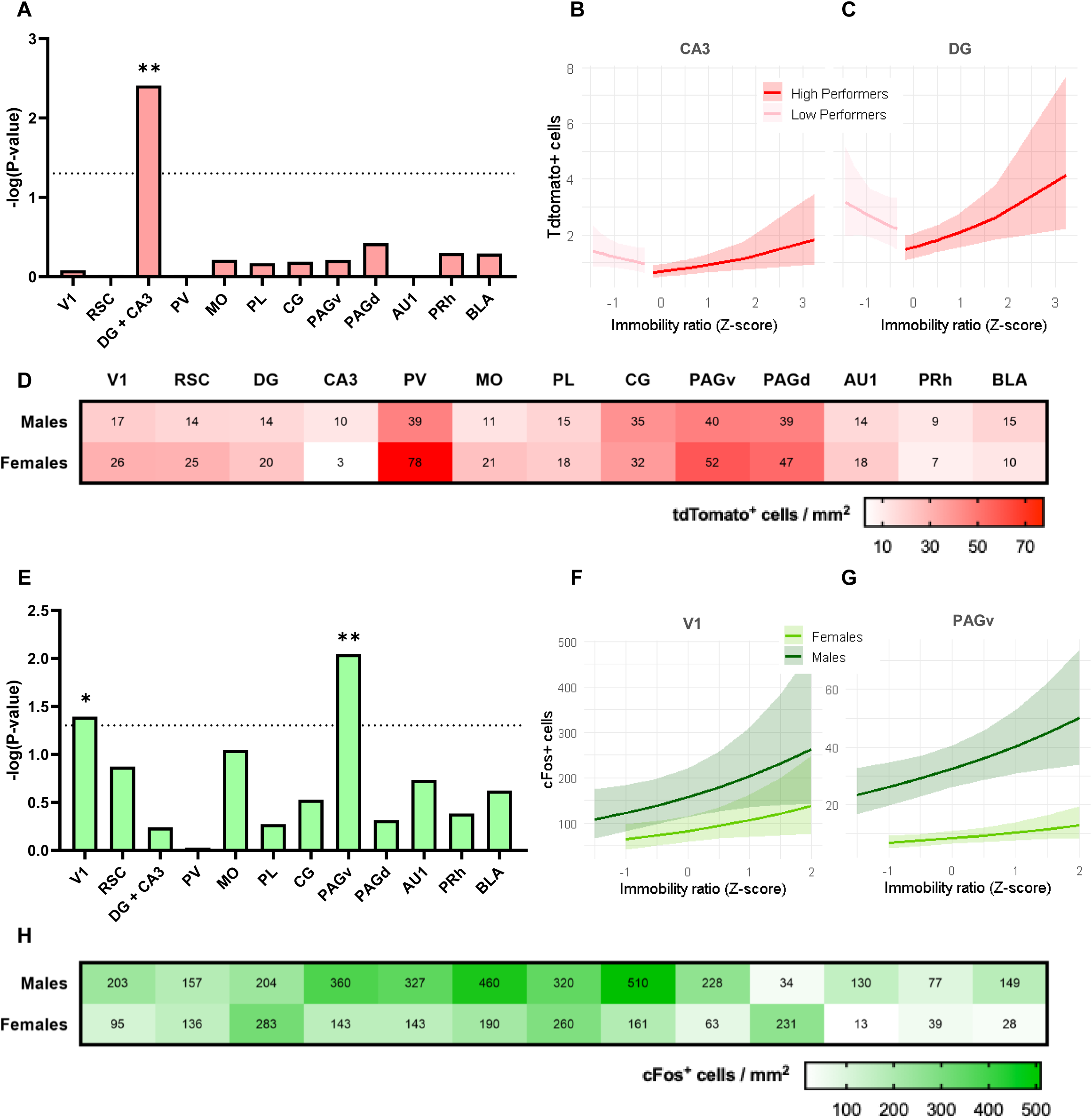
Quantification and modeling of tdTomato and cFos counts during memory recall. (A) Significance of the generalized linear mixed models (GLMM) coefficients of the tdTomato^+^ counts in each brain region by the immobility ratio: primary visual cortex (V1), retrosplenial cortex (RSC), dentate gyrus (DG), cornu ammonis 3 (CA3), paraventricular nucleus of the thalamus (PV), medial orbitofrontal area (MO), prelimbic cortex (PL), cingulate cortex (CG), ventral periaqueductal gray (PAGv), dorsal periaqueductal gray (PAGd), primary auditory area (AU1), perirhinal cortex (PRh), basolateral amygdala (BLA). Only the hippocampal DG and CA3 showed a significant coefficient (p = 0.00389, n = 23) for the CS_2_ recall. (B-C) tdTomato^+^ cells in the CA3 and DG for each immobility ratio, separated by performance (p = 0.0429, n = 23). In DG and CA3, high-performing mice exhibited increased tdTomato^+^ cell activation with immobility, whereas low-performing mice showed an opposite relationship. (D) Heatmaps of mean density of tdTomato^+^ cells separated by sex and brain region. (E) Significance of the GLMM coefficients of the cFos^+^ counts in each brain region by the immobility ratio. The V1 (p = 0.0405, n = 23) and PAGv (p = 0.00902, n = 23) yielded significant coefficients. (F-G) cFos^+^ cells in the V1 and PAGv for each immobility ratio. Both areas exhibit a positive trend, with males having higher values than females in both regions (V1: p = 0.00793, n = 23; PAGv: p *<*0.0001, n = 23). (H) Heatmaps of mean density of cFos^+^ cells separated by sex and brain region. P-values are denoted as p *<*0.05 (*), p *<*0.01 (**), and p *<*0.001 (***).

To assess differences in labeling across brain regions, we normalized raw counts by the area of each region of interest (ROI) and calculated the average cell density per region for each animal. Overall, females exhibited similar or slightly higher tdTomato levels than males in most areas, with the most pronounced difference in the PV, where females showed nearly twice the density (78 cells/mm^2^) compared to males (39 cells/mm^2^). Additional regions with notable female-dependent increases included the RSC, V1, MO, PAGd, and PAGv. In contrast, only a few regions, such as the CG, displayed higher counts in males, although these effects were modest. Some areas, including the CA3 and BLA, showed consistently low tdTomato counts in both sexes (Figure 3D and S1).

Regarding the cFos effects, indicating the immediate-early gene expression after CS_1_ recall, the two brain regions that showed a significant effect of immobility ratios were V1 (p = 0.0405) and PAGv (p = 0.00902) (Figure 3E). In both areas, higher immobility ratios corresponded with increased expected numbers of cFos^+^ cells. Specifically, in V1, a one-unit increase in the immobility ratio led to a 29.1% increase in cFos^+^ cell counts according to the model (Figure 3F), while in PAGv, the same increase in the ratio corresponded to a 24.2% rise in cFos labeling, holding other variables constant (Figure 3G). Additionally, both regions showed a notable effect of sex: the model estimated that males had approximately 91.5% more cFos^+^ cells than females in V1 (p = 0.00793), and nearly four times more in PAGv (p *<*0.0001).

In cFos labeling, the sex differences showed an opposite pattern compared to tdTomato. Males generally had higher cFos counts across nearly all regions, especially in the MO (460 vs. 190), CG (510 vs. 161), and V1 (203 vs. 95). In contrast, females exhibited relatively higher cFos activation only in the DG and PAGd, indicating region-specific sex differences in stimulus-induced activation (Figure 3H and S2).

Overall, the linear models on CS_2_ memory recall showed that behavioral performance significantly shaped neural activation in the dorsal DG and CA3. In high performers, increased immobility corresponded to higher tdTomato^+^ counts, while in low performers, the same trend reversed. These findings suggest that successful memory recall of second-order conditioned memories may involve dorsal hippocampal circuits, specifically DG and CA3.

### Inhibiting DG/CA3 CaMKII^+^ cells impairs second-order conditioning

To gain a deeper understanding of the role of DG and CA3 in SOC, we aimed to identify the cells activated during CS_2_ recall, specifically whether the tdTomato^+^ cells were glutamatergic or GABAergic. To achieve this, we employed an immunohistochemical approach, relying on the colocalization of the tdTomato signal with antibodies against CaMKII, a marker of glutamatergic neurons, and GAD67, a marker of GABAergic neurons (Figure 4A).

**Figure 4.**
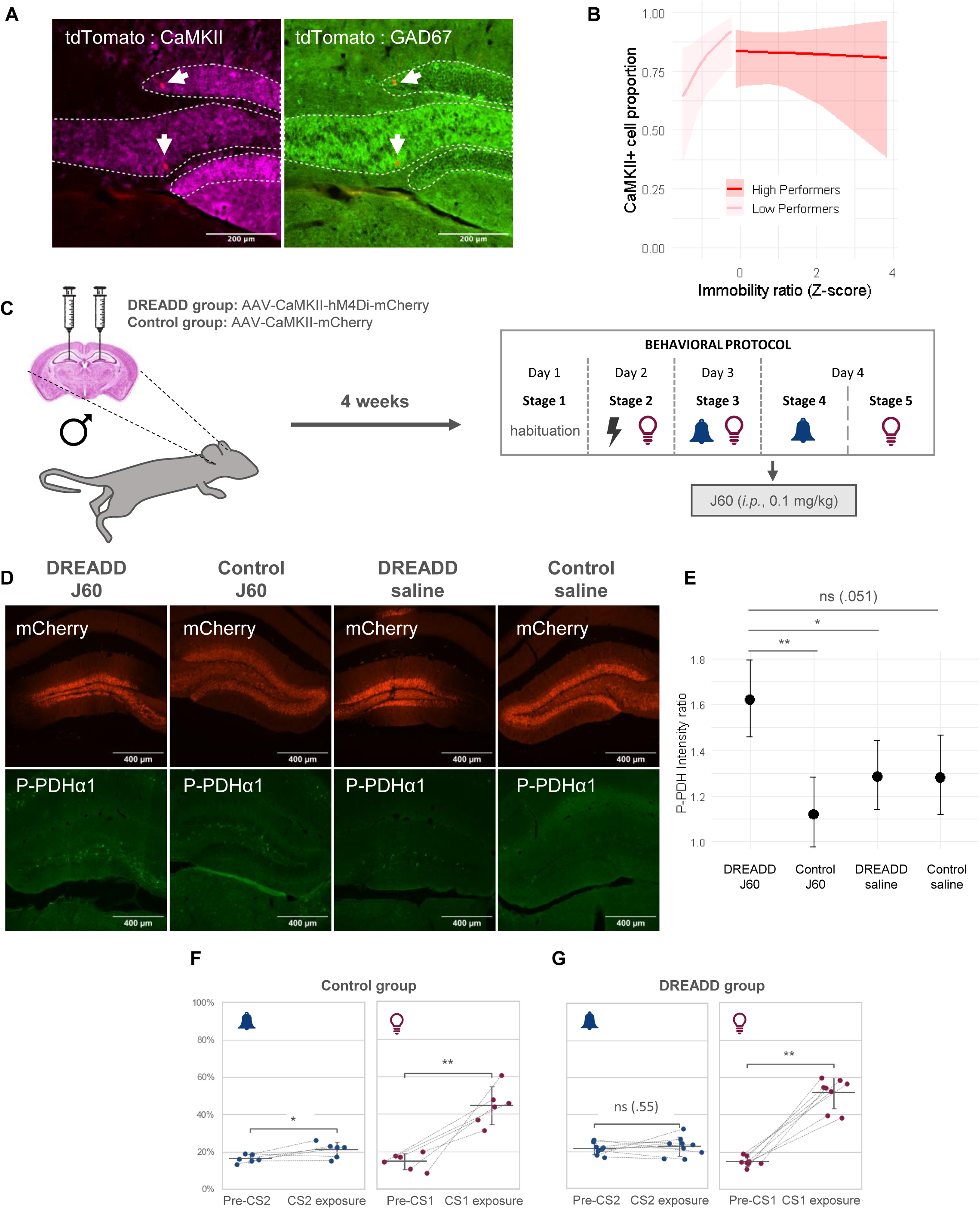
Role of CaMKII^+^ cells in DG/CA3 during SOC recall. (A) Immunohistochemical colocalization of tdTomato (red) with CaMKII^+^ cells (magenta) and GAD67^+^ cells (green) in DG and CA3, showing that most tdTomato^+^ cells are excitatory neurons expressing CaMKII. Dashed lines outline the hippocampal DG. Scale bars: 200 µm. (B) Proportion of CaMKII^+^ cells out of CaMKII^+^ and GAD67^+^ cells for each immobility ratio, separated by performance (p = 0.033, n = 23). High-performers maintain a stable CaMKII^+^ ratio of 70%, whereas low-performers show an increasing trend with immobility. (C) Experimental design schema. Male mice received bilateral DG/CA3 injections of AAV-CaMKII-hM4Di-mCherry (DREADD group) or control virus AAV-CaMKII-mCherry. After 4 weeks, mice underwent the behavioral SOC protocol, in which they received an injection of the DREADD agonist J60 (0.1 mg/kg, i.p.) 1 hour before CS_2_ recall testing. (D) Verification of viral expression: Representative images of mCherry fluorescence (top) indicating viral spread, and P-PDH*α*1 immunostaining (bottom) to measure neural inactivity. Scale bars: 400 µm. (E) Quantification of P-PDH*α*1 intensity ratio (DG/CA3 vs. surrounding tissue) shows significant inhibition of neural activity in the DREADD group treated with J60 compared to controls with J60 (p = 0.0030, n = 3), DREADD with saline (p = 0.0356, n = 4), and controls with saline (p = 0.0513, n = 3). (F-G) Behavioral effects of chemogenetic inhibition on immobility (huddle) during the SOC recall. The control group shows normal increases in immobility to both CS_1_ (p = 0.0443, n = 15) and CS_2_ (p = 0.0033, n = 15). The DREADD group exhibits an impaired immobility response to CS_2_ (p = 0.55, n = 15) but not to CS_1_ (p = 0.0078, n = 15). P-values are denoted as p *<*0.05 (*), p *<*0.01 (**), and p *<*0.001 (***). The data are presented as estimated intensity ratios with their 95% confidence interval on panel E, and mean ± SD on panels F and G.

We analyzed the results by fitting a GLMM under a binomial distribution, controlling for the same explanatory variables as the previous linear models. When examining the proportion of CaMKII^+^ cells across experimental conditions, we observed no significant effects for sex (p = 0.731) or between brain regions (i.e., DG and CA3, p = 0.210). However, group-specific effects of the immobility ratio on CaMKII^+^ proportions showed a significant interaction (p = 0.033). Among low performers, each one-unit increase in the immobility ratio resulted in a 4.37-fold increase in the expected proportion of CaMKII^+^ cells. This sharp rise, however, caused a saturation effect as the response ratio continued to increase. High performers maintained a relatively stable CaMKII^+^ proportion of 80% across the entire range of immobility ratios, with no significant modulation (Figure 4B). Therefore, this result suggests that the majority of labeled cells are excitatory neurons.

By combining the CaMKII^+^ cell proportions with the tdTomato^+^ counts, we suggest that CaMKII^+^ cells in the dorsal hippocampus, particularly within the DG/CA3 regions, may contribute to modulating CS_2_ memory recall. To test its causal involvement, we employed a chemogenetic approach using designer receptors exclusively activated by designer drugs (DREADDs). We bilaterally injected male mice in the dorsal hippocampus, spreading accross DG/CA3 with either AAV-CaMKII-hM4Di-mCherry (hereafter, DREADD group) or AAV-CaMKII-mCherry (hereafter, control group). This chemogenetic approach enabled selective inhibition of CaMKII^+^ cells in the targeted hippocampal regions during the retrieval of CS_1_ and CS_2_ memories. After four weeks to ensure adequate viral expression, the animals underwent the standard behavioral protocol used to induce and measure first- and second-order conditioned responses. On the recall day, mice received a 0.1 mg/kg intraperitoneal injection of the DREADD agonist JHU37160 dihydrochloride (J60)^41^ one hour before the CS_2_ exposure test (Figure 4C).

To confirm proper viral expression in the targeted cells, we assessed the localization of the viral constructs in both DREADD and control groups (Figure 4D). In addition, to validate the DREADD-dependent inhibition of hippocampal circuits, we divided each group into two subgroups: one received a saline injection, and the other received J60 to modulate the DREADD receptors (i.e., inhibiting cell activity). One hour after injection, we perfused the mice and processed the brain tissue to immunolabel Phospho-Pyruvate Dehydrogenase (P-PDH*α*1), which inversely correlates with neural activity^24^. cFos expression typically reflects neural activation and may be less reliable for indicating decreases in activity following DREADD-mediated inhibition. To quantify changes in neural inhibition, we calculated the ratio of P-PDH*α*1 signal between the targeted (DG/CA3) and surrounding brain tissue. We analyzed this metric using a linear mixedeffects model (LMM) with fixed effects for brain region, viral construct, injection type, and their interaction. We included mouse identity as a random intercept to account for repeated measures across areas. The model showed that mice expressing the inhibitory DREADD and injected with J60 had a significantly higher P-PDH*α*1 intensity ratio, about 1.6 times, indicating decreased neural activity in the targeted areas. In contrast, the same mice injected with saline had a significantly lower ratio of only 1.2 (p = 0.0356). The control groups with J60 and saline also displayed baseline activity, with ratios of 1.1 and 1.3, respectively (Figure 4E). These results confirm the functional expression of the viral constructs and the effectiveness of the chemogenetic manipulation in CaMKII^+^ hippocampal neurons (Figure 4D).

After validating our chemogenetic approach, we evaluated behavioral responses in both DREADD and control groups using huddling, which is the same immobility measure that showed significant modulation in response to CS_1_ and CS_2_ in TRAP2:Ai14 mice. The control group exhibited significant behavioral changes in response to both CS_2_ (p = 0.0443) and CS_1_ (p = 0.0033) (Figure 4F). In contrast, in the DREADD group, inhibition of DG/CA3 selectively disrupted the response to CS_2_ (p = 0.55), while the response to CS_1_ remained unaffected (p = 0.0078) (Figure 4G). Overall, these findings suggest that CaMKII^+^ cells in the DG/CA3 are necessary for the recall of second-order, but not first-order, conditioned memories.

## DISCUSSION

By combining pose tracking, genetic tagging, automated image analysis, and chemogenetic manipulation, this study investigates the brain substrates underlying the expression of SOC and FOC in a light-tone SOC protocol in mice. Using TRAP2:Ai14 male and female mice, we permanently labeled neurons activated during memory retrieval in SOC and FOC and identified brain-wide activity patterns^29^. Behavioral analysis using DeepLabCut^31^ and Keypoint-MoSeq^32^ revealed consistent changes in syllable usage, particularly increased immobility, in response to both CS1 and CS2, establishing huddling^33^ as a reliable and straightforward behavioral marker for SOC recall. Modeling brain activity showed that CS_2_-induced behavioral responses could explain the activity of specific excitatory neurons (CaMKII^+^ cells) in the DG/CA3 regions of the dorsal hippocampus. Notably, the chemogenetic inhibition of these cells impaired the recall of SOC memories without affecting the recall of FOC, underscoring the crucial role of hippocampal excitatory neurons in the expression of SOC.

The data-driven analysis of TRAP2:Ai14 animals confirms our previous findings on SOC behavioral responses in wild-type C57BL/6J mice^30^: (1) behavioral responses to CS_1_ and CS_2_ in our SOC paradigm remain unaffected by sex, (2) responses to CS_2_ are generally less intense than those to CS_1_, and, most importantly, (3) CS_1_ and CS_2_ evoke intrinsically distinct behav-ioral patterns. Rather than simply mirroring first-order behavioral responses, SOC elicits unique behavioral profiles for each conditioned stimulus. In particular, as the syllable analysis shows, responses to CS_1_ mainly consist of immobility, whereas CS_2_ evokes a more diverse behavioral repertoire, including both passive behaviors, such as immobility, and active responses, such as exploration and escape-like movements. However, to correlate brain activity alongside behavioral performance, we relied on immobility (i.e., huddling) as our primary metric. Immobility emerged as the most robust and reliable behavioral indicator of associative memory during SOC, with mean Δ values between 3 and 14 times higher than other behavioral syllables. Although linear models that include the full syllable quantification might achieve higher coefficients of determination, we deliberately prioritized interpretability over marginal improvements in model fit. Including all syllables causes severe multicollinearity, which weakens the stability and reliability of linear estimates^42^. Even when applying techniques to address this problem, such as dimensionality reduction and regularization, models with multiple features are more complex to biologically interpret. Conversely, one metric offers a clear, meaningful, and reproducible measure that we can strongly associate with neural activity^43,44^. By focusing on this well-defined feature, we maintain interpretable and actionable results, particularly for posterior manipulations (e.g., chemogenetic approaches), where a solid behavioral anchor is essential for assessing causal effects^45^.

After having established a behavioral metric, we measured the pattern of neural activity in SOC (tdTomato^+^) and FOC (cFos^+^), which displayed sex-dependent differences. tdTomato labeling during CS_2_ recall was generally higher in female mice, whereas cFos expression following CS_1_ recall was more pronounced in males. These differences may reflect sex-specific strategies or patterns of brain circuit engagement during recall, in line with prior reports indicating that female rodents often recruit broader neural networks for associative learning^46^. Nonetheless, we cannot entirely exclude methodological confounds, such as sex differences in 4-OHT metabolism^47^ or baseline expression of immediate-early genes^48,49^. Future studies incorporating additional controls may determine whether these differences reflect true biological dimorphism or arise from technical variability.

Regardless of sex, we found that certain areas of the brain were strongly activated during CS_2_ recall. These regions include the PV, PAGv, and PAGd, which are structures commonly linked to emotional and survival-related processes. The PV is involved in arousal, attention, and stress regulation^50–52^, indicating that SOC recall may engage circuits related to emotional salience and autonomic control, even without a direct pairing with the US. Similarly, the PAG, known for its role in fear, defensive behavior, and pain modulation^53,54^, showed strong activation, supporting the idea that CS_2_ can trigger internal fear-related states. Future experiments will interrogate the potential causal involvement of these brain regions, which strongly activate during light-tone associations, in SOC formation.

During first-order memory recall (CS_1_), cFos activation in the PAGv and V1 significantly corresponded to better behavioral performance, specifically immobility. The PAGv is classically associated with freezing and passive defensive responses to aversive stimuli^7,55,56^, while activation in V1 may reflect enhanced visual sensory input during light presentation^56,57^. These findings support the idea that CS_1_ recall engages established defensive and sensory circuits.

In contrast, during SOC recall, tdTomato expression in the DG and CA3 significantly corresponded to greater immobility responses. Notably, the DG facilitates pattern separation, transforming similar inputs into distinct representations, while CA3 supports pattern completion, enabling the retrieval of complete memories from partial or ambiguous cues^58–60^. Therefore, these findings suggest that DG and CA3 activity may help facilitate the inference of threat from indirect associations, such as CS_2_, which was never directly paired with the US. Importantly, chemogenetic inhibition confirmed a causal role for excitatory neurons in these hippocampal regions during the expression of CS_2_ responses, while leaving CS_1_ unaffected. This difference indicates that dorsal hippocampal excitatory activity is necessary for recalling second-order, but not first-order, associations. While CS_1_ memory relies on a direct US pairing, typically supported by subcortical circuits such as the amygdala, PAG, and hypothalamus^7,8^, our SOC paradigm seems to require an additional inference step (i.e., linking CS_2_ to CS_1_, and then to the US^5,61,62^).

In this context, hippocampal excitatory neurons may mediate complex associative learning, enabling animals to infer threat from stimuli that are only indirectly linked to the US^58–60^. Thus, while DG and CA3 activity may be dispensable for direct associations (FOC), it appears to be critical for higher-order associations (SOC). Previous studies support this interpretation, at least in the context of sensory preconditioning (SPC). For example, the dorsal hippocampus is necessary to encode associations between two neutral stimuli during preconditioning, but not for learning a neutral cue association with the US^16^. Similarly, type 1 cannabinoid receptors (CB1R) in hippocampal GABAergic neurons are both necessary and sufficient for SPC, but not for direct conditioning^63^. Although some studies have also implicated the dorsal hippocampus in FOC with light or tone stimuli^64–66^, the overall body of evidence supports the idea that higher-order conditioning engages dorsal hippocampal mechanisms specifically tuned to inferring relationships between indirectly associated stimuli.

Throughout our study, we analyzed tdTomato and cFos expression using our self-developed CellRake package, a scalable and accurate Python tool for the automated detection of fluorescently labeled cells. We benchmarked CellRake by training separate models for tdTomato and cFos. Although both classifiers performed well, the tdTomato model consistently outperformed the cFos model across all metrics, including computing speed. This difference likely reflects the intrinsic properties of the markers: cFos, a nuclear marker, produces dimmer, smaller, and often overlapping signals, while tdTomato, a cytoplasmic marker, generates larger, more uniformly labeled cells with higher contrast against the background. Indeed, the nature of each fluorescent marker can also explain these differences, since tdTomato is triggered by genetic recombination, whereas cFos depends heavily on antibody-based labeling. However, despite these differences, CellRake achieved strong agreement with manual counts of both markers, and its design could enable easy adaptation to other markers or experimental conditions. This tool also offers several advantages over existing software. Unlike broader platforms such as QuPath^67^, CellProfiler^68^, Ilastik^69^, or Cellpose^28^, which provide extensive but often complex functionalities, CellRake is designed for a single task, resulting in a distribution size of under 30 KB and a significantly lower learning curve. Its intuitive design allows both advanced users and beginners to work efficiently from day one. The annotation process is fast and precise, relying on Python tools without the need for external programs like ImageJ^70^, thereby eliminating time-consuming manual steps such as drawing regions of interest around blobs to train the model. Additionally, CellRake runs efficiently on standard hardware by using scikit-learn models^37^ rather than GPU-intensive deep learning methods, making it a more accessible and practical choice for researchers who lack high-performance computing resources.

Taken together, our findings identify excitatory neurons in the DG/CA3 as key substrates supporting SOC expression. Future research should investigate how hippocampal excitatory and inhibitory circuits interact during the different stages of SOC aversive paradigms and whether these mechanisms apply to other types of second-order associations (e.g., using appetitive cues) or how they may become dysregulated in models of psychiatric disorders. Overall, by combining advanced behavioral measurement, temporal cell tagging, scalable histological analysis, and targeted circuit inhibition, this study advances our understanding of how the brain learns from indirect experiences.

## METHODS

### Animals

We used male and female TRAP2:Ai14 mice for the identification of brain regions involved in the recall of SOC, aged 8-10 weeks, genetically engineered to carry two crucial elements: Fos-CreERT and the Ai14 reporter cassette. These animals came from a cross between two homozygous strains: TRAP2 mice (Fos^tm2.1(icre/ERT2)Luo/^J; The Jackson Laboratory, USA; #030323) and Ai14 tdTomato reporter mice (B6.Cg-Gt(ROSA)26Sor^tm14(CAG–tdTomato)Hze^/J; The Jackson Laboratory; #007914).

For inhibition of CaMKII^+^ cells in the dorsal hippocampus, we used male C57BL/6J mice, aged 8-10 weeks, obtained from Charles River Laboratories (Spain).

We housed all mice in groups of four males or five females, never mixing both sexes. We kept the animals in an inverted 12-hour light cycle (lights turned off at 7:30 a.m.) and conducted the experiments in darkness under red light. We provided food and water *ad libitum* and controlled the temperature (22 ± 1°C) and humidity (50 ± 5%). The appropriate animal care and use committee of our institution and the Generalitat de Catalunya approved all the procedures stated.

### Second-order conditioning protocol

We used the same light-tone behavioral protocol for mouse light-tone second-order conditioning (SOC) described and validated in our previous work^30^. Briefly, it consisted of five stages: habituation to the conditioning chambers (day 1), pairing a light (CS_1_) with an electric shock (day 2), pairing the previously conditioned CS_1_ with a novel tone (CS_2_) (day 3), and an independent probe test for the CS_2_ and the CS_1_ (day 4). Using this behavioral protocol, which involves observing specific CS_1_- and CS_2_-mediated behavioral responses, we can study the brain substrates of SOC.

### Pose estimation and behavioral analysis

We followed the methodological framework described in our previous work^30^. Briefly, the procedure involved recording videos of mice during the recall stages and analyzing them using DeepLabCut 3.0^31^, with a focus on 14 body parts. To identify the behavioral structure, we applied Keypoint-MoSeq^32^, an unsupervised clustering tool that captures the frequency and timing of behavioral motifs. In parallel, we used the supervised DeepOF pipeline^33^ to extract huddling behavior, which acts as a proxy for fear-related responses^30^.

We divided the animals into high- and low-performing groups based on their huddling responses to CS_1_ and CS_2_. We calculated a response ratio by dividing the immobility percentage during the stimulus period by that of the preceding baseline period. To define the threshold, we selected the lowest response ratio observed for CS_1_, which was 1.5. Animals with a response ratio below 1.5 were classified as low performers, while those with ratios above this threshold were considered high performers.

### Drug preparation and administration

We prepared 4-hydroxytamoxifen (4-OHT; Merck, #H6278) at 20 mg/mL in ethanol by gently shaking it at 37°C for 5 minutes. We then aliquoted the solution and stored it at –20°C. Before use, we mixed 4-OHT with Kolliphor oil (Merck, Cat #C5135-500G) by shaking at 37°C for 5 minutes. We remove ethanol using a Savant DNA120 SpeedVac concentrator, which results in a concentration of 10 mg/mL. Finally, we added 0.1 M phosphate-buffered saline to adjust the solution to a final dose of 25 mg/kg. We immediately administered the 4-OHT by intraperitoneal injection just after assessing the mice’s response to the tone (CS_2_).

We prepared JHU37160 dihydrochloride (J60; Hello Bio, #HB6261) in sterile saline to achieve a final concentration of 0.1 mg/kg. We administered J60 or saline intraperitoneally 1 hour before the probe test, during which CS_2_ exposure occurred.

### Induction of tdTomato expression in TRAP2:Ai14

Immediately after assessing the mice’s response to the tone (CS_2_), we injected a solution of 25 mg/mL 4-hydroxytamoxifen (4-OHT) intraperitoneally at a ratio of 0.1 mg/kg. The flow of 4-OHT induces the expression of the tdTomato marker protein in the cells activated during the CS_2_-recalling session. We then waited one week before assessing the recall of the light (CS_1_) to allow enough time for tdTomato to express itself. Then, we evaluated the response to the light, and 90 minutes after the stimulus was presented, we prepared the mice for perfusion.

### Perfusion and sectioning of brains

One week after 4-OHT administration and CS_2_ exposure, we assessed the recall of the light (CS_1_) to allow enough time for tdTomato to express itself. We prepared the mice for perfusion 90 minutes after displaying the light stimulus. Following a modified protocol from^71^, we transcardiacally perfused the mice. Briefly, we anesthetized them using a ketamine (100 mg/kg) and xylazine (10 mg/kg) mixture and then perfused them with a 4% paraformaldehyde (PFA) solution in 0.1 M PB, pH 7.4. We stored the brains individually in 4% PFA-filled tubes at 4°C and, after 24 hours, transferred them to a 30% sucrose solution for cryoprotection at the same temperature. After three days, we froze the brains and cut coronal sections at a thickness of 30 µm using a Leica CM 1950 cryostat at -20°C. We stored the slices in cryoprotectant (30% ethylene glycol and 20% glycerol in 0.02 M PB) until their usage.

### Immunofluorescence staining

We followed an immunohistological protocol as previously described^72,73^ to detect cFos, CaMKII, GAD67, and Phospho-Pyruvate Dehydrogenase (P-PDH*α*1) in perfused brain slices. We rinsed the collected brain sections three times with 0.1 M phosphate buffer (PB) for 5 minutes each. The tissue was then permeabilized with a 0.2% Triton X-100 solution (in PB) for five 5-minute intervals and blocked with 5% normal donkey serum (NDS) for one hour at room temperature. After blocking non-specific bindings, we incubated the slices overnight at 4°C with the primary antibody at a 1:1000 dilution in the remaining blocking solution. The primary antibodies were rabbit anti-cFos (SynapticSystem, #226008), rabbit anti-CaMKII (Abcam, #EP1829Y), mouse anti-GAD67 (Sigma Aldrich, #MAB5406), or rabbit anti-P-PDH*α*1 (Cell Signaling Technology, #37115).

The next day, we rinsed the slices again with a 0.2% Triton X-100 solution three times, followed by a 2-hour incubation at room temperature in the dark with the secondary antibody solution at a 1:1000 dilution. The secondary antibodies used were donkey anti-rabbit AlexaFluor 488-conjugated (Jackson ImmunoResearch, #711-545-152) or donkey anti-mouse AlexaFluor 647-conjugated (Jackson ImmunoResearch, #712-605-150).

After incubation, we rinsed the sections five times with a 0.5% Triton X-100 solution, air-dried them on microscope slides, and mounted them with Fluoromount-G containing DAPI for nuclear staining. We stored the slides at 4°C and protected them from light.

### Imaging of brain slices

We imaged the stained brain sections using a Nikon Eclipse Ni-E fluorescence microscope with NIS-Elements software. At 10X magnification, we captured four to six images of each animal’s region of interest in the brain. In TRAP2:Ai14 mice, we selected 13 brain regions: primary visual cortex (V1, Bregma = -3.64), retrosplenial cortex (RSC, Bregma = -1.82), dentate gyrus (DG, Bregma = -1.82), cornu ammonis 3 (CA3, Bregma = -1.82), paraventricular nucleus of the thalamus (PV, Bregma = -1.82), medial orbitofrontal area (MO, Bregma = 2.46), prelimbic cortex (PL, Bregma = 1.70), cingulate cortex (CG, Bregma = -0.22), ventral periaqueductal gray (PAGv, Bregma = -4.60), dorsal periaqueductal gray (PAGd, Bregma = -4.60), primary auditory area (AU1, Bregma = -3.64), perirhinal cortex (PRh, Bregma = -1.82), basolateral amygdala (BLA, Bregma = -1.82).

### CellRake’s segmentation algorithm

CellRake begins the image analysis by processing each grayscale fluorescence image with a multistep segmentation pipeline. The purpose of segmentation is to identify well-defined regions of interest (ROIs) that correspond to positive signal detections. First, it detects potential blob-like structures using a Laplacian of Gaussian (LoG) filter^74^, which enhances local maxima in intensity and captures structures of varying sizes. These detections serve as seeds for further processing. CellRake extracts a circular area centered on each detection for every candidate region and applies Otsu’s method^75^ to perform adaptive thresholding around the specific blob. This localized approach accounts for background intensity heterogeneity. Following thresholding, CellRake uses morphological operations to eliminate small holes within blobs and discard irregular or fragmented detections based on compactness and area constraints. The algorithm then applies a marker-based watershed algorithm to separate overlapping signals. Finally, CellRake transforms the final segmented shapes into polygonal contours, clips them to remain within image bounds, and stores them in a structured dictionary format for downstream classification.

### CellRake’s classification algorithm

We implemented a semi-supervised learning pipeline based on label spreading to classify segmented blobs. First, CellRake extracts 22 morphological and intensity-based features for each blob identified during the segmentation step. These features serve as input to a k-means clustering algorithm (with a default of 10 clusters) to facilitate stratified manual labeling. The user manually labels 10 representative blobs from each cluster as positive or negative.

Given the typical class imbalance, which has fewer positives, we apply the Synthetic Minority Oversampling Technique (SMOTE) to oversample the minority class. CellRake then performs label spreading using a K-Nearest Neighbors (KNN) kernel to propagate labels across the dataset. To ensure high-confidence labels, we compute the entropy of the posterior class probabilities for each blob and retain only those predictions with low entropy (less than 0.025). CellRake then uses these confidently labeled samples to perform a train-test split for supervised model training and evaluation. Finally, it outputs five performance metrics: area under the receiver operating characteristic curve (ROC AUC), average precision (AP), precision, recall, and F1 score.

### CellRake’s colocalization of markers

Identifying colocalizations (i.e., markers that overlap significantly in the same image) is often valuable in experimental designs with multiple markers. CellRake supports this by using images that were previously analyzed for each marker individually. After independent analyses, CellRake calculates the intersection of all the detected markers in each image. In this case, it considers that two markers colocalize if the overlapped area constitutes at least 80% of the smaller area.

### Stereotaxic surgeries

We anesthetized mice via intraperitoneal injection of ketamine (75 mg/kg) combined with medetomidine (1 mg/kg). We provided analgesia through a subcutaneous injection of meloxicam (5 mg/kg) administered before surgery and for two consecutive days postoperatively. Once anesthetized and after confirming the absence of a paw pinch reflex, we secured the animal’s head in a stereotaxic frame and aligned it using bregma and lambda coordinates. We then made two small craniotomies to allow for bilateral infusion of the viral vector solution into the DG/CA3 from the dorsal hippocampus. The stereotaxic coordinates used were: anteroposterior (AP) -2 mm, mediolateral (ML) *±*1.5 mm, and dorsoventral (DV) -2.1 mm.

We injected 500 nL of either AAV-CaMKII-hM4Di-mCherry (Addgene viral prep #50477-AAV2, titer *≥*7 *×* 1,012 vg/mL) or AAV-CaMKII-mCherry (Addgene viral prep #114469-AAV5, titer *≥*7 *×* 1,012 vg/mL). After the injections, we sutured the incision site and allowed the mice to recover for 3 to 4 weeks before initiating the behavioral protocol.

### Statistical methods

To analyze the behavioral data, we used the same approach described in our recent work^30^. In brief, for each behavioral syllable obtained from Keypoint-MoSeq^32^, we calculated the difference between the time spent before and after stimulus presentation, known as Δ values. To assess whether syllable usage changed significantly after cue presentation, we applied Hotelling’s T^2^ test on the multivariate distribution of Δ scores per sex and stimulus condition. We further analyzed behavioral dynamics through transition matrices, where we computed Manhattan distances (MD) between pre- and post-stimulus periods. We assessed statistical significance using bootstrap-based empirical p-values.

To assess significant differences between conditions in continuous univariate variables, we used two-tailed t-tests (paired or unpaired, depending on the design), as well as one-way or two-way ANOVA for comparisons involving multiple groups. We verified normality assumptions using the Shapiro–Wilk test and homogeneity of variances using Levene’s test. When normality was violated, we applied non-parametric alternatives, such as the Wilcoxon signed-rank test or Mann–Whitney U test, as appropriate.

To validate CellRake, we compared automated counts against manual annotations. We first assessed bivariate normality through the Henze-Zirkler multivariate normality test. We used Pearson’s correlation for normally distributed data and Spearman’s correlation otherwise.

To test the relationship between brain activation and behavior, we fitted generalized linear mixed models (GLMMs) separately for each brain region. We modeled cell counts (cFos or tdTomato) as the response variable with log-transformed ROI area as an offset, and animal ID as a random intercept to account for repeated measures. Fixed effects included sex, behavioral performance group, and immobility ratio, with interaction terms when appropriate. Initially, we assumed a Poisson distribution; however, due to overdispersion, we refitted the models with a Negative Binomial distribution. For the tdTomato data, which displayed excess zeros, we tested multiple zero-inflated models and selected the best-fitting one based on the Akaike Information Criterion (AIC).

To determine the cellular identity of activated neurons, we used binomial GLMMs modeling the probability of tdTomato^+^ cells being CaMKII^+^ across brain regions and performance groups. Fixed effects included immobility ratio, sex, and their interaction, with animal ID as a random effect.

To confirm effective neural silencing via DREADDs, we analyzed P-PDH*α*1 intensity ratios (targeted vs. surrounding tissue) using a linear mixed-effects model (LMM) with fixed effects for region, viral construct, injection type, and their interactions, as well as mouse identity as a random intercept.

All statistical comparisons used the following significance thresholds: p *<*0.05 (*), p *<*0.01 (**), and p *<*0.001 (***).

## Supporting information

Statistical tables

## RESOURCE AVAILABILITY

### Lead contact

Requests for further information and resources should be directed to and will be fulfilled by the lead contact, Arnau Busquets-Garcia, PhD.

### Materials availability

This study did not generate new materials.

### Data and code availability

- Any additional information required to reanalyze the data reported in this paper is available from the lead contact upon request.

## ACKNOWLEDGMENTS

We want to thank the personnel of the Animal Facility of the Parc de Recerca Biomedica de Barcelona (PRBB) for mouse care. We thank all the members of our lab for valuable discussions during the development of the project. This work was supported by the Generalitat de Catalunya (SGR-00022) from the Departament d’Economia i Coneixement de la Generalitat de Catalunya (Spain) and from the European Research Council (ERC) under the European Union’s Horizon 2020 research and innovation program (Grant agreement Num. [948217]). The project that gave rise to these results received the support of a fellowship from “la Caixa” Foundation (ID 100010434). The fellowship code is LCF/BQ/DR22/11950014.

## AUTHOR CONTRIBUTIONS

A.B-G. and M.C. contributed to the conception of the project. M.C., J.B-A., D.R-C., and J.G-P performed and analyzed all experiments. J.P. helped in setting up the SOC protocol. A.B-G. and M.C. wrote and revised the manuscript. All authors approved the final version of the manuscript.

