## Supplementary Figures for "Dentate gyrus and CA3 activity mediates light-tone second-order conditioning expression in mice"

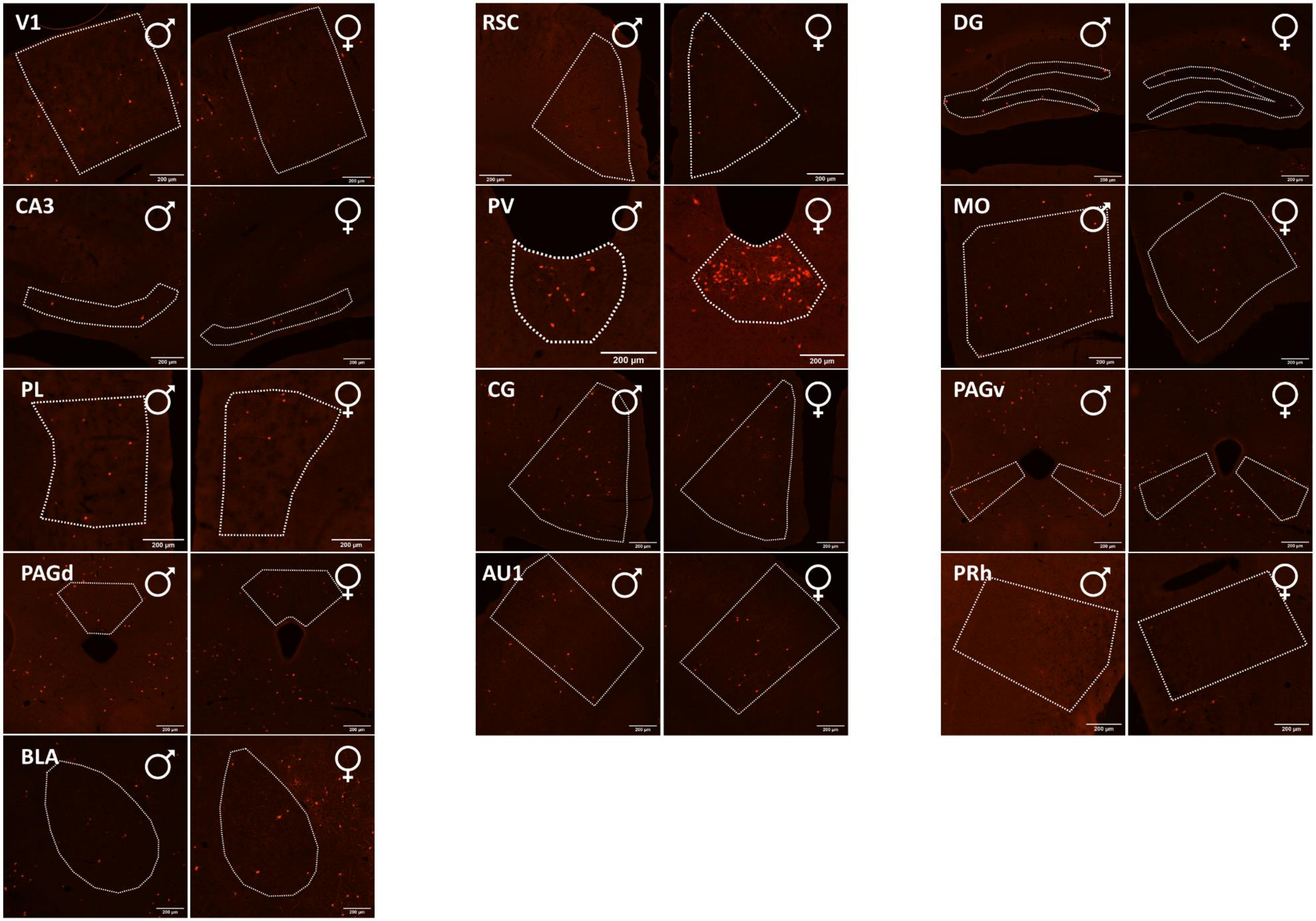


**Figure S1. Representative images for all 13 brain areas of both sexes for the tdTomato marker.** The analyzed brain areas are the primary visual cortex (V1), retrosplenial cortex (RSC), dentate gyrus (DG), cornu ammonis 3 (CA3), paraventricular nucleus of the thalamus (PV), medial orbitofrontal area (MO), prelimbic cortex (PL), cingulate cortex (CG), ventral periaqueductal gray (PAGv), dorsal periaqueductal gray (PAGd), primary auditory area (AU1), perirhinal cortex (PRh), and basolateral amygdala (BLA). The white dotted line indicates the contour of each region. Scale bars: 200 μm.

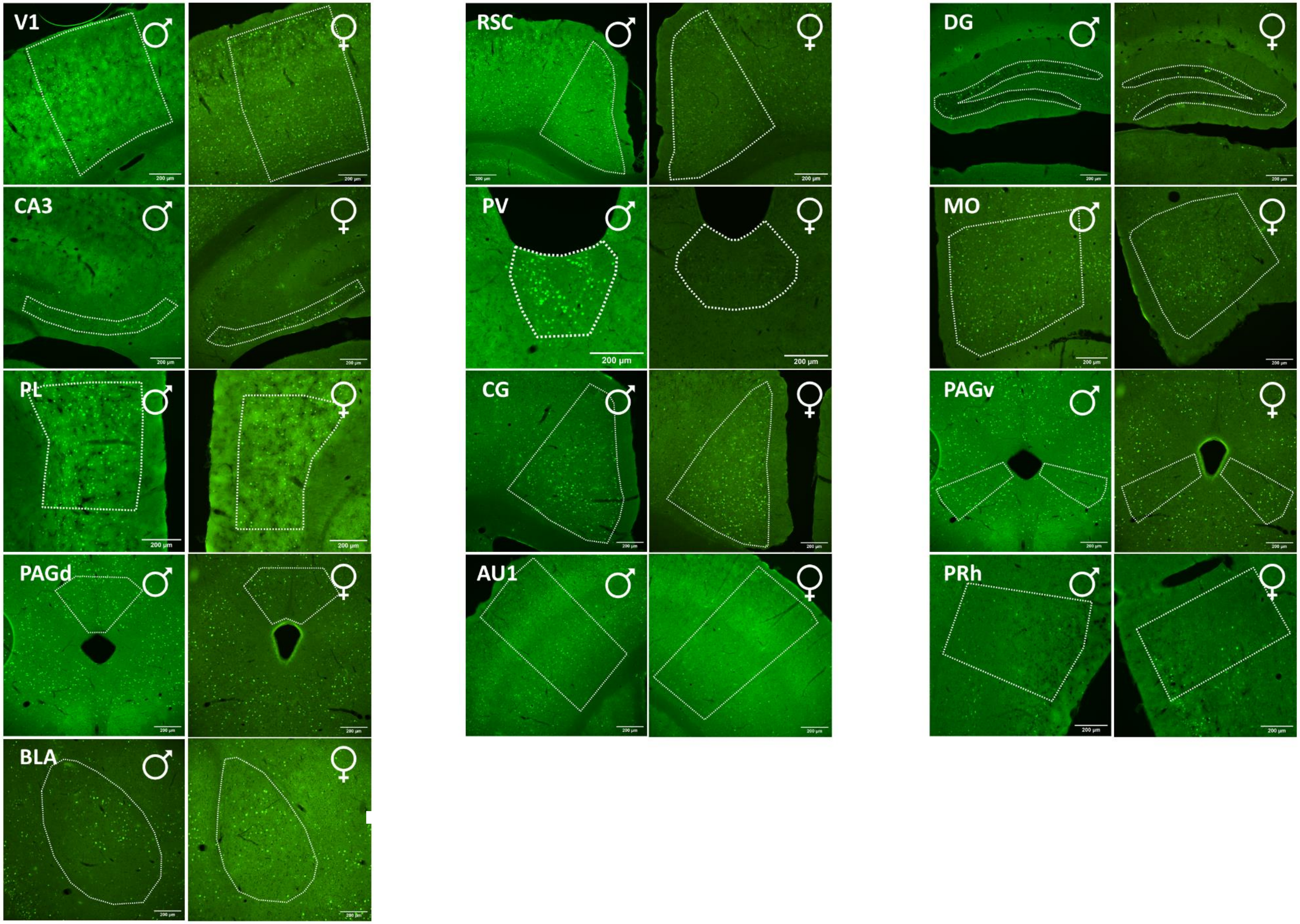


**Figure S2. Representative images for all 13 brain areas of both sexes for the cFos marker.** The analyzed brain areas are the primary visual cortex (V1), retrosplenial cortex (RSC), dentate gyrus (DG), cornu ammonis 3 (CA3), paraventricular nucleus of the thalamus (PV), medial orbitofrontal area (MO), prelimbic cortex (PL), cingulate cortex (CG), ventral periaqueductal gray (PAGv), dorsal periaqueductal gray (PAGd), primary auditory area (AU1), perirhinal cortex (PRh), and basolateral amygdala (BLA). The white dotted line indicates the contour of each region. Scale bars: 200 μm.
