## Supplementary material for "Dentate gyrus and CA3 activity mediates light-tone second-order conditioning expression in mice": Statistical tables

| Figure | Experiment | Size | Normality | Statistical test | P- value |
|---|---|---|---|---|---|
| Figure 1 | (B) $CS_1$ males | 12 | Yes | Hotelling's $T^2$ test | p = 4.219e-6 |
| | (B) $CS_2$ males | 12 | Yes | Hotelling's $T^2$ test | p = 0.0210 |
| | (C) $CS_1$ females | 11 | Yes | Hotelling's $T^2$ test | p = 4.869e-6 |
| | (C) $CS_2$ females | 11 | Yes | Hotelling's $T^2$ test | p = 0.0167 |
| | (D) $CS_1$ males | 12 | - | Permutation test | p = 1.99e-8 |
| | (D) $CS_2$ males | 12 | - | Permutation test | p = 1.14e-4 |
| | (E) $CS_1$ females | 11 | - | Permutation test | p = 1.62e-4 |
| | (E) $CS_2$ females | 11 | - | Permutation test | p = 0.0424 |
| | (G) Syllable 0 $CS_1$ males vs females | 11<br>12 | Yes | Two-tailed t-test | p = 0.616 |
| | (G) Syllable 0 $CS_2$ males vs females | 11<br>12 | Yes | Two-tailed t-test | p = 0.999 |
| | (H) Syllable 2 $CS_1$ males vs females | 11<br>12 | Yes | Two-tailed t-test | p = 0.999 |
| | (H) Syllable 2 $CS_2$ males vs females | 11<br>12 | Yes | Two-tailed t-test | p = 0.997 |
| | (I) $CS_2$ males | 12 | No | Wilcoxon signed-rank test | p = 0.00195 |
| | (I) $CS_1$ males | 12 | No | Wilcoxon signed-rank test | p = 0.0005 |
| | (J) $CS_2$ females | 11 | Yes | Paired t-test | p = 0.0167 |
| | (J) $CS_1$ females | 11 | Yes | Paired t-test | p < 0.0001 |
| Figure 2 | (E) Eccentricity | 990<br>12642 | No | Mann-Whitney U Test | p = 0.5447 |
| | (E) Area | 990<br>12642 | No | Mann-Whitney U Test | p < 0.0001 |
| | (E) Perimeter | 990<br>12642 | No | Mann-Whitney U Test | p < 0.0001 |
| | (E) Intensity ratio | 990<br>12642 | No | Mann-Whitney U Test | p < 0.0001 |
| | (E) Intensity difference | 990<br>12642 | No | Mann-Whitney U Test | p < 0.0001 |
| | (E) Mean intensity | 990<br>12642 | No | Mann-Whitney U Test | p < 0.0001 |
| | (E) SD intensity | 990<br>12642 | No | Mann-Whitney U Test | p < 0.0001 |
| | (H) Correlation cFos | 153 | No | Spearman's correlation | r = 0.9440<br>p < 0.0001 |
| | (I) Correlation tdTomato | 153 | No | Spearman's correlation | r = 0.8789<br>p < 0.0001 |
| | (J) Correlation colocalization | 153 | Yes | Pearson's correlation | r = 0.9124<br>p < 0.0001 |
| Figure 3 | (C) GLMM tdTomato V1 | 23 | - | Negative binomial GLMM | p = 0.829 |
| | (C) GLMM tdTomato RSC | 23 | - | Negative binomial GLMM | p = 0.948 |
| | (C) GLMM tdTomato DG + CA3 | 23 | - | Negative binomial GLMM | p = 0.00389 |
| | (C) GLMM tdTomato PV | 23 | - | Negative binomial GLMM | p = 0.948 |
| | (C) GLMM tdTomato MO | 23 | - | Negative binomial GLMM | p = 0.611 |
| | (C) GLMM tdTomato PL | 23 | - | Negative binomial GLMM | p = 0.676 |
| | (C) GLMM tdTomato CG | 23 | - | Negative binomial GLMM | p = 0.648 |
| | (C) GLMM tdTomato PAGv | 23 | - | Negative binomial GLMM | p = 0.616 |
| | (C) GLMM tdTomato PAGd | 23 | - | Negative binomial GLMM | p = 0.379 |

| | | | | | |
|---|---|---|---|---|---|
| | (C) GLMM tdTomato AU1 | 23 | - | Negative binomial GLMM | p = 0.971 |
| | (C) GLMM tdTomato PRh | 23 | - | Negative binomial GLMM | p = 0.503 |
| | (C) GLMM tdTomato BLA | 23 | - | Negative binomial GLMM | p = 0.511 |
| | (F) GLMM cFos V1 | 23 | - | Negative binomial GLMM | p = 0.0405 |
| | (F) GLMM cFos RSC | 23 | - | Negative binomial GLMM | p = 0.134 |
| | (F) GLMM cFos DG + CA3 | 23 | - | Negative binomial GLMM | p = 0.577 |
| | (F) GLMM cFos PV | 23 | - | Negative binomial GLMM | p = 0.934 |
| | (F) GLMM cFos MO | 23 | - | Negative binomial GLMM | p = 0.0898 |
| | (F) GLMM cFos PL | 23 | - | Negative binomial GLMM | p = 0.535 |
| | (F) GLMM cFos CG | 23 | - | Negative binomial GLMM | p = 0.297 |
| | (F) GLMM cFos PAGv | 23 | - | Negative binomial GLMM | p = 0.00902 |
| | (F) GLMM cFos PAGd | 23 | - | Negative binomial GLMM | p = 0.484 |
| | (F) GLMM cFos AU1 | 23 | - | Negative binomial GLMM | p = 0.184 |
| | (F) GLMM cFos PRh | 23 | - | Negative binomial GLMM | p = 0.412 |
| | (F) GLMM cFos BLA | 23 | - | Negative binomial GLMM | p = 0.239 |
| Figure 4 | (E) DREADD J60 vs Control J60 | 15 | - | LMM | p = 0.0030 |
| | (E) DREADD J60 vs DREADD saline | 15 | - | LMM | p = 0.0356 |
| | (E) DREADD J60 vs Control saline | 15 | - | LMM | p = 0.0513 |
| | (F) DREADD $CS_2$ | 9 | Yes | Paired t-test | p = 0.558 |
| | (F) DREADD $CS_1$ | 9 | No | Wilcoxon signed-rank test | p = 0.00781 |
| | (G) Control $CS_2$ | 6 | Yes | Paired t-test | p = 0.0443 |
| | (G) Control $CS_1$ | 6 | Yes | Paired t-test | p = 0.00326 |

**Table S1.** Statistical analysis related to Main Figures 1-4.
